# RGS10 deficiency facilitates distant metastasis by inducing epithelial-mesenchymal transition in breast cancer

**DOI:** 10.1101/2024.03.04.583283

**Authors:** Yang Liu, Yi Jiang, Peng Qiu, Tie Ma, Yang Bai, Jiawen Bu, Yueting Hu, Ming Jin, Tong Zhu, Xi Gu

## Abstract

Distant metastasis is the major cause of death in patients with breast cancer. Epithelial–mesenchymal transition (EMT) contributes to breast cancer metastasis. Regulator of G protein-signaling (RGS) proteins modulate metastasis in various cancers. This study identified a novel role for RGS10 in EMT and metastasis in breast cancer. RGS10 protein levels were significantly lower in breast cancer tissues compared to normal breast tissues, and deficiency in RGS10 protein predicted a worse prognosis in patients with breast cancer. RGS10 protein levels were lower in the highly aggressive cell line MDA-MB-231 than in the poorly aggressive, less invasive cell lines MCF7 and SKBR3. Silencing RGS10 in SKBR3 cells enhanced EMT and caused SKBR3 cell migration and invasion. The ability of RGS10 to suppress EMT and metastasis in breast cancer was dependent on lipocalin-2 and miR-539-5p. These findings identify RGS10 as a tumor suppressor, prognostic biomarker, and potential therapeutic target for breast cancer.

## Introduction

Breast cancer is the most common cancer among women worldwide. Globally, in 2020, an estimated 2.3 million new cases of breast cancer were diagnosed, and there were approximately 685,000 deaths from the disease (Sung et al., 2021). The majority of breast cancer mortality is due to distant metastasis, with 5-year survival estimated at 30% (American Cancer Society, 2023). There is a critical need to identify early breast cancer metastasis using prognostic biomarkers to ensure patients receive effective anticancer therapies in a timely manner (Miglietta et al., 2022).

Epithelial-to-mesenchymal transition (EMT) plays a critical role in tumor progression and metastatic invasion in breast cancer. EMT describes the process by which epithelial cells lose their epithelial characteristics and cell-cell contact and increase their invasive potential (Singh and Chakrabarti, 2019). EMT is characterized by a loss of epithelial cell markers, such as cytokeratins and E-cadherin, followed by an upregulation in the expression of mesenchymal cell markers, such as N-cadherin and vimentin. EMT is regulated at different levels by factors involved in cell signaling, transcriptional control, and epigenetic and post-translational modifications (Lai et al., 2020).

Biomarkers of EMT may predict early breast cancer metastasis and facilitate clinical decision-making (Liu et al., 2016). The regulator of G protein signaling 10 (RGS10) belongs to the superfamily of RGS proteins that bind and deactivate heterotrimeric G proteins (Almutairi et al., 2020). RGS proteins are important mediators of essential cellular processes and may be tumor initiators or tumor suppressors (Li et al., 2023). The canonical function of RGS proteins is to act as GTPase-activating proteins (GAPs), accelerate GTP hydrolysis on G-protein alpha subunits, and terminate signaling pathways downstream of G protein-coupled receptors (Hunt et al., 1996). RGS proteins can have noncanonical GAP-independent functions, including the suppression of transforming growth factor beta (TGF-β)-induced EMT in non-small cell lung cancer by RGS6 (Wang et al., 2022).

RGS10 is a critical regulator of cell survival, polarization, adhesion, chemotaxis, and differentiation that exhibits tumor-suppressing effects in ovarian and colorectal cancer. In ovarian cancer cells, RGS10 suppression increases proliferation by phosphorylation of mTOR, 4E-BP1, p70S6K and rProtein-S6, including in the presence of chemotherapy (Altman et al., 2015), loss of RGS10 expression contributes to the development of chemoresistance (Cacan et al., 2014), and modulating RGS10 expression can alter sensitivity to paclitaxel, cisplatin, and vincristine (Cacan et al., 2014; Hooks et al., 2010; Hooks and Murph, 2015). In ovarian tumors, transcription of RGS10 is regulated by DNA methylation and histone deacetylation (Nguyen et al., 2014). In colorectal cancer, RGS10 expression is suppressed, and inhibition of DNA methylation may contribute to improved prognosis (Caldiran and Cacan, 2022).

In breast cancer, RGS10 is downregulated in heavily metastatic human breast cancer cell populations compared to weakly metastatic human breast cancer cell populations (Montel et al., 2005; Steeg, 2005). The mechanisms underlying the metastasis-suppressing function of RGS10 in breast cancer remain to be elucidated. In this study, we investigated the function of RGS10 in breast cancer, specifically in breast cancer metastasis. The objectives were to 1) characterize the expression of RGS10 in freshly resected breast cancer and adjacent normal breast tissues; 2) determine the prognostic significance of RGS10 expression in patients with breast cancer; and 3) explore the role of RGS10 and upstream effectors in tumor progression and metastasis in breast cancer cells *in vitro* and *in vivo*.

## Methods

### Key resources table

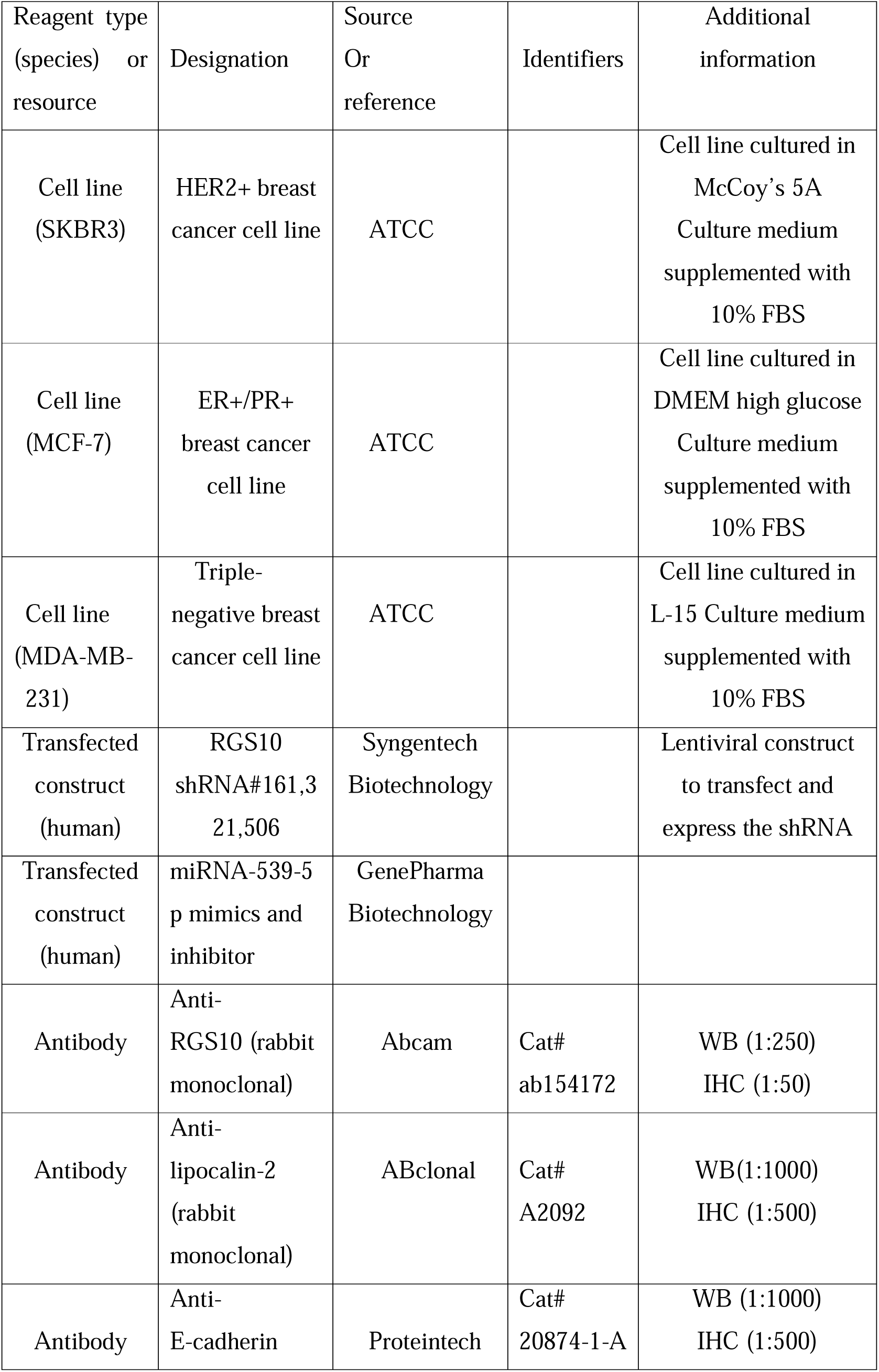
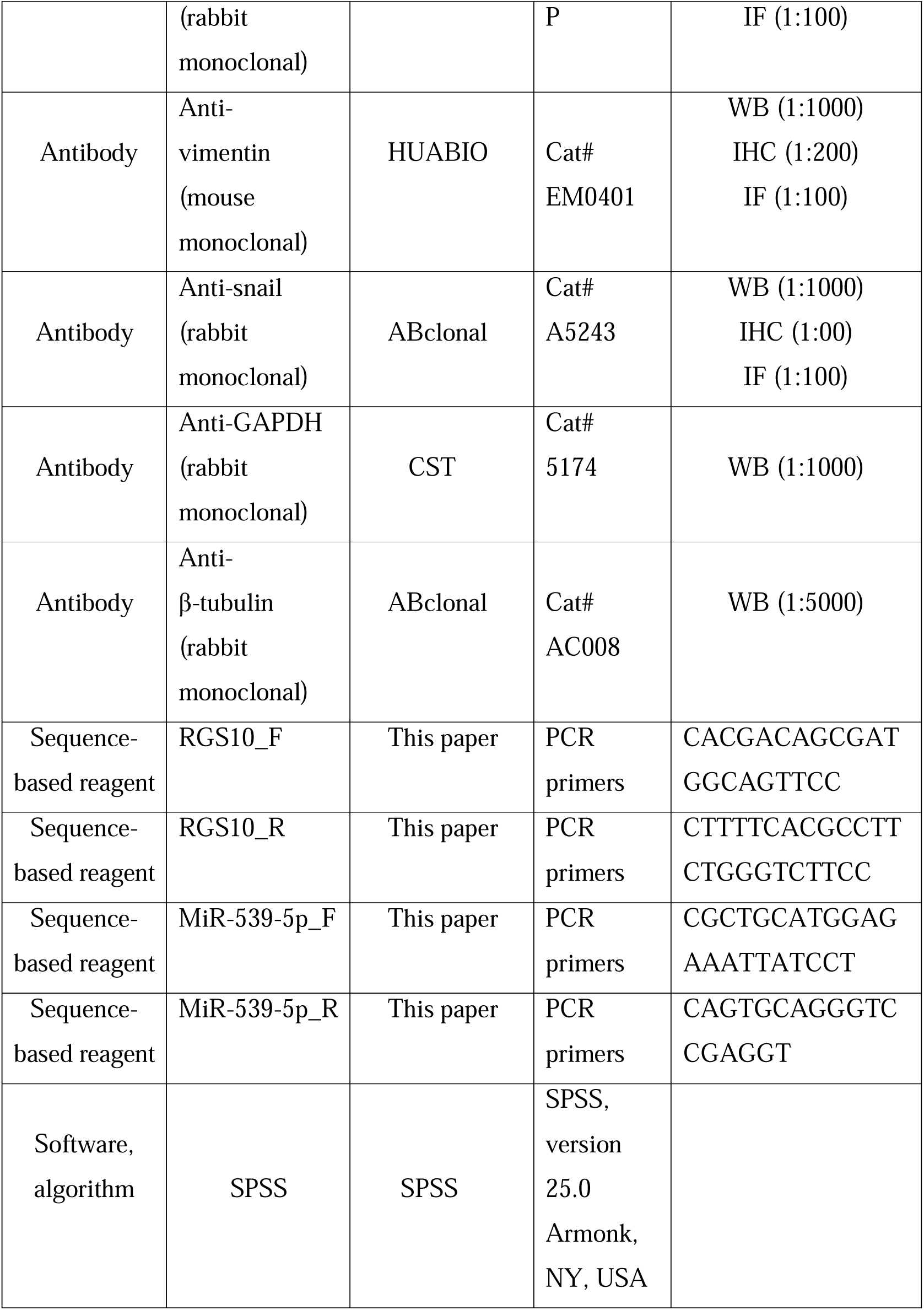

### Bioinformatics

The Genotype-*Tissue Expression* database (www.gtexportal.org) was used to analyze RGS10 mRNA levels in 31 normal human tissues. The Cancer Cell Line Encyclopedia database (https://sites.broadinstitute.org/ccle) was used to analyze RGS10 mRNA levels in a series of cell lines (Rahman et al., 2019). The Kaplan–Meier plotter (http://kmplot.com) was used to assess the relevance of RGS10 mRNA expression for disease-free survival (DFS) and overall survival (OS) in patients with breast cancer (Győrffy, 2021).

### Clinical specimens

This study included an additional 153 patients with histologically confirmed invasive ductal breast carcinoma who received treatment at the Shengjing Hospital of China Medical University from April 2006 to April 2008. Patients had: 1) undergone surgery with no distant metastases at the time of the operation; 2) received standard adjuvant therapy after surgery; 3) ≥10 axillary lymph nodes dissected and examined pathologically after surgery (Coates et al., 2015; Gradishar et al., 2021; Wolff et al., 2013); and 4) ≥10 years of follow-up as outpatients or by telephone interviews.

Twenty paired breast cancer and normal adjacent tissues were obtained from patients undergoing surgical resection at Shengjing Hospital of China Medical University to evaluate the status of RGS10. Clinical specimens from 133 patients were used to assess the effect of RGS10 on prognosis. The clinicopathological characteristics of these 153 patients are shown in Table 1 and Table 2.

**Table 1.** Clinicopathological characteristics of 20 patients.

| Clinicopathological characteristic | Patient 1 | Patient 2 | Patient 3 | Patient 4 | Patient 5 |
| --- | --- | --- | --- | --- | --- |
| Sex | Female | Female | Female | Female | Female |
| Age (years) | 58 | 40 | 70 | 38 | 58 |
| T grade | 2 | 1 | 2 | 1 | 1 |
| N grade | 0 | 1 | 3 | 0 | 1 |
| Menopausal status | Postmenopausal | Premenopausal | Postmenopausal | Premenopausal | Postmenopausal |
| Subclass | Luminal A | Luminal A | Luminal A | Luminal A | Luminal A |
| Metastasis | No | No | No | No | No |
|  | Patient 6 | Patient 7 | Patient 8 | Patient 9 | Patient 10 |
| Sex | Female | Female | Female | Female | Female |
| Age (years) | 59 | 56 | 80 | 55 | 37 |
| T grade | 2 | 2 | 3 | 2 | 1 |
| N grade | 3 | 0 | 2 | 2 | 3 |
| Menopausal status | Postmenopausal | Postmenopausal | Postmenopausal | Postmenopausal | Premenopausal |
| Subclass | Luminal B | Luminal B | Luminal B | Luminal B | Luminal B |
| Metastasis | No | No | No | No | No |
|  | Patient 11 | Patient 12 | Patient 13 | Patient 14 | Patient 15 |
| Sex | Female | Female | Female | Female | Female |
| Age (years) | 62 | 57 | 41 | 53 | 58 |
| T grade | 2 | 2 | 2 | 1 | 2 |
| N grade | 3 | 1 | 3 | 3 | 1 |
| Menopausal status | Postmenopausal | Postmenopausal | Premenopausal | Postmenopausal | Postmenopausal |
| Subclass | HER2 positive | HER2 positive | HER2 positive | HER2 positive | HER2 positive |
| Metastasis | No | No | No | No | No |
|  | Patient 16 | Patient 17 | Patient 18 | Patient 19 | Patient 20 |
| Sex | Female | Female | Female | Female | Female |
| Age (years) | 50 | 37 | 35 | 51 | 55 |
| T grade | 2 | 2 | 2 | 2 | 2 |
| N grade | 0 | 0 | 0 | 0 | 0 |
| Menopausal status | Premenopausal | Premenopausal | Premenopausal | Premenopausal | Postmenopausal |
| Subclasses | Triple negative | Triple negative | Triple negative | Triple negative | Triple negative |
| Metastasis | No | No | No | No | No |

HER2, human epidermal growth factor receptor 2.

**Table 2.** Correlations between RGS10 expression and clinicopathological characteristics.

| Variable | RGS10 expression<br>(n, %) | RGS10 expression (n, %) | P-value |
| --- | --- | --- | --- |
| No. of patients | 34 (25.6) | 99 (74.4) |  |
| Age (years) |  |  | 0.590 |
| ≤45 | 8 (23.5) | 28 (28.3) |  |
| >45 | 26 (76.5) | 71 (71.7) |  |
| Other disease |  |  | 0.746 |
| Yes | 28 (82.4) | 79 (79.8) |  |
| No | 6 (17.6) | 20 (20.2) |  |
| Histological grade |  |  | 0.744 |
| I | 12 (35.3) | 30 (30.3) |  |
| II | 19 (55.9) | 56 (56.6) |  |
| III | 3 (8.8) | 13 (13.1) |  |
| Tumor size (cm) |  |  | 0.101 |
| Median (range) | 2.191 (1.0–8.0) | 1.861 (0.6–4.0) |  |
| No. of positive axillary lymph nodes |  |  | 0.316 |
| Median (range) | 1.06 (0–19) | 1.93 (0–27) |  |
| No. of pregnancies |  |  | 0.706 |
| Median (range) | 2.50 (1–6) | 2.40 (0–7) |  |
| No. of births |  |  | 0.497 |
| Median (range) | 1.62 (1–3) | 1.75 (0–7) |  |
| Age at first pregnancy (years) |  |  | 0.489 |
| <21 | 1 (2.9) | 6 (6.1) |  |
| 21–25 | 15 (44.1) | 42 (42.4) |  |
| 26–30 | 18 (52.9) | 46 (46.5) |  |
| >30 | 0 (0) | 5 (5.0) |  |
| Menopausal status |  |  | 0.216 |
| Premenopausal | 21 (61.8) | 49 (49.5) |  |
| Postmenopausal | 13 (38.2) | 50 (50.5) |  |
| Estrogen receptor status |  |  | 0.397 |
| Positive | 25 (73.5) | 65 (65.7) |  |
| Negative | 9 (26.5) | 34 (34.3) |  |
| Progesterone receptor status |  |  | 0.527 |
| Positive | 25 (73.5) | 78 (78.8) |  |
| Negative | 9 (26.5) | 21 (21.2) |  |
| HER2 status |  |  | 0.496 |
| Positive | 6 (17.6) | 23 (23.2) |  |
| Negative | 28 (82.4) | 76 (76.8) |  |
| Ki67 status |  |  | 0.773 |
| >20 | 14 (41.2) | 38 (38.4) |  |
| ≤20 | 20 (58.8) | 61 (61.6) |  |
| Molecular Type |  |  | 0.043 |
| Luminal A | 6 (17.6) | 30 (30.3) |  |
| Luminal B | 19 (55.9) | 36 (36.3) |  |
| HER2+ | 2 (5.9) | 16 (16.2) |  |
| Triple-negative breast cancer | 7 (20.6) | 17 (17.2) |  |
| Distant metastasis |  |  | 0.008 |
| Yes | 7 (20.6) | 46 (46.5) |  |
| No | 27 (79.4) | 53 (53.5) |  |
| Disease-free survival<br>(months) |  |  | <0.001 |
| Median (range) | 114.71 (53–140) | 89.66 (9–132) |  |
| Death |  |  | 0.024 |
| Yes | 5 (14.7) | 35 (35.4) |  |
| No | 29 (85.3) | 64 (64.6) |  |
| Overall survival (months) |  |  | 0.030 |
| Median (range) | 120.41 (80–140) | 107.64 (24–140) |  |

HER2, human epidermal growth factor receptor 2.

The study was approved by the Institutional Ethics Committee of Shengjing Hospital of China Medical University and complied with the principles of the Declaration of Helsinki and Good Clinical Practice guidelines of the National Medical Products Administration of China. Informed consent was obtained from all the participants.

### Breast cancer cells and culture

Human breast cancer cell lines SKBR3, MCF-7, and MDA-MB-231 were obtained from American Type Culture Collection (ATCC, Manassas, VA). The identity of cell lines had been authenticated by using STR profiling and with no contamination in the mycoplasma test. SKBR3 cells were cultured in McCoy’s 5A (modified) medium supplemented with 10% fetal bovine serum at 37°C and 5% CO_2_. MCF7 cells were cultured in DMEM high glucose (Invitrogen) medium supplemented with 10% fetal bovine serum at 37°C and 5% CO_2_. MDA-MB-231 cells were cultured in Leibovitz’s L-15 medium supplemented with 10% fetal bovine serum at 37°C without CO_2_.

### Transfection

SKBR3 cells were transfected with short hairpin (sh)RNAs specifically targeting RGS10 (Syngentech Biotechnology, Beijing, China). shRNA sequences were: shRNA-RGS10-161, 5′-GCCTCAAGAGCACAGCCAAAT-3′; shRNA-RGS10-321, 5′-GGAGATCTACATGACCTTTCT-3′; a n d shRNA-RGS10-506, 5′-GCACCCTCTGATGTTCCAGAA-3′. The shRNA-negative control (NC) sequence was 5′-AAACGTGACACGTTCGGAGAA-3′. As transfection efficiencies of shRNA-RGS10-161 and shRNA-RGS10-506 were better than shRNA-RGS10-321, shRNA-RGS10-161 and shRNA-RGS10-506 were used for subsequent experiments.

A miR-539-5p mimic, miR-539-5p inhibitor, and corresponding NC were designed and synthesized by GenePharma Biotechnology (Shanghai, China). The sequences of miRNA were: miR-539-5p mimic, 5′-GGAGAAAUUAUCCUUGGUGUGU-3′; miR-539-5p mimic NC,5′-UUCUCCGAACGUGUCACGUTT-3′; miR-539-5p inhibitor,5′-ACACACCAAGGAUAAUUUCUCC-3′; miR-539-5p inhibitor NC 5′-CAGUACUUUUGUGUAGUACAA′. SKBR3 cells were transfected with the miR-539-5p mimic or NC. MDA-MB-231 cells were transfected with the miR-539-5p inhibitor or NC.

Lipofectamine 3000 (L3000015, Thermo Fisher Scientific, Invitrogen, USA) was used for cell transfection.

### Reverse transcription-quantitative polymerase chain reaction (RT-qPCR)

Total RNA was isolated from breast cancer tissues and breast cancer cell lines using Triquick reagent (Solarbio Life Science, Beijing, China) and reverse-transcribed using a cDNA synthesis kit (Takara Bio, Kusatsu, Japan). RT-qPCR was conducted using SYBR Green PCR Master Mix (TaKaRa) on a Real-time PCR System. Primers were: RGS10 forward, 5′-CACGACAGCGATGGCAGTTCC-3′; RGS10 reverse, 5′-CTTTTCACGCCTTCTGGGTCTTCC-3′; GAPDH forward, 5′-CCTTCCGTGTCCCCACT-3′; and GAPDH reverse, 5′-GCCTGCTTCACCACCTTC-3′. Thermocycling conditions were 95°C for 30 s, 40 cycles of 95°C for 3 s, and 60°C for 30 s. Relative mRNA levels were calculated using the comparative threshold (Cq) cycle method (2^-ΔΔCt^) (Adnan et al., 2011).

cDNA synthesis in SKBR3 cells transfected with the miR-539-5p mimic or NC and MDA-MB-231 cells transfected with the miR-539-5p inhibitor or NC was conducted using a miRNA First Strand cDNA Synthesis kit that adopted A-tailing (B532461, Sangon Biotech). qPCR was performed using a miRNA qPCR kit (B532461, Sangon Biotech). Relative miR-539-5p level was normalized to U6 as the internal control. Primers were as follows: miR-539-5p forward, 5′-CGCTGCATGGAGAAATTATCCT-3′; miR-539-5p reverse, 5′-CAGTGCAGGGTCCGAGGT-3′; U6 forward, 5′-GCTCGCTTCGGCAGCACATATAC-3′, and U6 reverse 5′-CGAATTTGCGTGTCATCCTTGCG-3′. Thermocycling conditions were 95°C for 30 s, 40 cycles of 95°C for 5 s, and 60°C for 30 s. Relative mRNA levels were calculated using the 2^-ΔΔCt^ method.

### Cell Counting Kit-8 (CCK-8) analysis

Cells were seeded into 96-well plates at a density of 5000 cells per well. After 0, 24, 48, and 72 h, CCK-8 (Cofitt Life Science Biotechnology) solution was added to each well, and cells were incubated for 3 h. Optical density was measured at 450 nm using a microplate spectrophotometer (Bio-Rad). Experiments were performed in triplicate.

### Colony formation assay

SKBR3 and MDA-MB-231 cells were trypsinized and plated on 6-well plates at a density of 1000 cells per well. When visible colonies were formed, the cells were fixed with methyl alcohol and stained with crystal violet. The plates were photographed, and the number of colonies was counted using ImageJ software.

### Cell invasion and migration assay

For cell invasion, 20,000 cells in 0.2 mL serum-free culture medium were placed in the upper chamber of a Transwell chamber that had been precoated with Matrigel matrix. The lower chamber was filled with 1 mL culture medium containing 10% fetal bovine serum as the chemoattractant. After culturing at 37°C for 24 h, cells that had migrated to the lower surface through the membrane were fixed with 4% paraformaldehyde for 10 min. Fixed cells were stained with crystal violet.

Detection of cell migration was similar to cell invasion, except the upper chamber of the Transwell chamber was not precoated with Matrigel.

### Immunohistochemistry

Surgically resected tissues were fixed in 4% formaldehyde, embedded in paraffin, and sectioned into 6-μm-thick slices. Sections were deparaffinized with xylene, rehydrated in a series of graded ethanols, and rinsed in Tris-buffered saline. Sections were incubated with a primary antibody anti-RGS10 (1:50; Abcam, Cambridge, MA; ab154172), anti-lipocalin-2 (LCN2) (1:500, ABclonal, #A2092), anti-E-cadherin (1:500, Proteintech, #20874-1-AP), anti-vimentin (1:200, HUABIO, #EM0401), anti-snail (1:100, ABclonal, # A5243), at 4°C overnight, incubated with a secondary antibody (Zhong Shan Jin Qiao Biotechnology Co., Beijing, China) at room temperature for approximately 45 min, and protein was visualized with a diaminobenzidine staining kit (Zhong Shan Jin Qiao Biotechnology Co.).

### Immunofluorescence staining

After transfection with the miR-539-5p mimic or miR-539-5p inhibitor for 24 h, SKBR3, and MDA-MB-231 cells were smeared on glass slides, cultured for 36 h, and fixed with 4% paraformaldehyde for 20 min. Cells were washed three times with phosphate-buffered saline (PBS) and blocked with 5% bovine serum albumin in PBS (pH 7.4) at room temperature for 1 h. Cells were incubated with anti-E-cadherin (1:100, Proteintech, #20874-1-AP), anti-vimentin ( 1:100, HUABIO, #EM0401), or anti-snail (1:100, ABclonal, #A5243) antibody at 4°C overnight. Cells were washed three times with PBS, and incubated with Alexa-Fluor 594 goat anti-mouse IgG H&L or Alexa-Fluor 488 goat anti-rabbit IgG H&L (Proteintech, 1:400 dilution) at room temperature for 1 h in the dark, washed three times, and incubated with 4′,6-diamidino-2-phenylindole (Solarbio, #C0065) for 10 min. After staining, cells were sealed with mounting media. Pictures were taken under a fluorescence microscope (Nikon, Japan).

### Western blotting

Cells were harvested by trypsinization and lysed in radioimmunoprecipitation assay lysis buffer. The cell lysate was centrifuged to a pellet (insoluble material). Proteins in the supernatant (30 µg of protein/lane) were separated by sodium dodecyl sulfate-polyacrylamide gel electrophoresis and transferred to polyvinylidene fluoride membranes. After blocking with 5% w/v nonfat dry milk for 1 h, membranes were incubated with anti-RGS10 (1:250; Abcam; ab154172), anti-GAPDH (1:1000, CST, 5174S), anti-lipocalin-2 (LCN2) (1:1000, ABclonal, #A2092), anti-E-cadherin (1:1000, Proteintech, #20874-1-AP), anti-vimentin (1:1000, HUABIO, #EM0401), anti-snail (1:1000, ABclonal, # A5243), or anti-β-tubulin (loading control) (1:5000, ABclonal, AC008) antibody at 4°C overnight. Membranes were incubated with a secondary antibody (Zhong Shan Jin Qiao Biotechnology Co.) at 37°C for 1 h. Bands were visualized using an enhanced chemiluminescent reagent. The relative protein levels of RGS10, LCN2, E-cadherin, vimentin, and snail were determined by densitometric analysis using Image Lab^TM^ 6.0.1 software.

### Dual-luciferase reporter assay

The StarBase database was used to predict potential binding sites between RGS10 mRNA and hsa-miR-539-5p. Sequences containing the 3′-untranslated region fragments of RGS10 and the mutated binding site of hsa-miR-539-5p or wild type were designed and constructed by GeneChem Biotechnology (Shanghai, China). The miR-539-5p mimic or NC was cotransfected with the wild-type or mutant plasmid using lipofectamine 3000 (Invitrogen). Luciferase intensity was recorded 48 h after transfection using the dual-luciferase reporter assay system (Promega, Madison, WI).

### Molecular interaction networks

The Search Tool for the Retrieval of Interacting *Genes*/Proteins database (https://string-db.org/) was used to identify proteins that interact with RGS10 and conduct a protein-protein network interaction analysis (Szklarczyk et al., 2015). Subsequently, the Database for Annotation, Visualization, and Integrated Discovery (https://david.ncifcrf.gov) was used to perform Gene Ontology (GO) and Kyoto Encyclopedia of Genes and Genomes (KEGG) pathway-enrichment analysis.

### *In-vivo* animal experiments

Nine BALB/C female nude mice (6 weeks old; 19–22 g) (Beijing Huafukang Biotechnology Company) were randomly assigned into three groups (n = 5 each). Mice were maintained in a specific-pathogen-free environment at 28°C and 50% humidity. The animal experiments were manipulated by the Regulations for the Administration of Affairs Concerning Experimental Animals and were approved by the Experimental Animal Ethics Committee of the Shengjing Hospital of China Medical University. A total of 1 × 10^7^ SKBR3 cells transfected with shRNA-RGS10 or shRNA-NC were injected subcutaneously to induce tumors. Tumor volume was measured every 3 days and calculated as V=1/2 (width^2^ × length). After subcutaneous injection of tumor cells for 30 days, mice were humanely euthanized and tumors were dissected and analyzed.

### Statistical analysis

Statistical analyses were conducted using SPSS 25.0 software. *In-vitro* experiments were performed in triplicate. Associations between the RGS10 gene and protein expression and clinicopathological characteristics were evaluated with the chi-squared test or independent samples *t*-test, as appropriate. Survival curves were generated using the Kaplan-Meier method. Univariate and multivariate Cox regression analyses were used to identify prognostic predictors related to DFS and OS. Hazard ratios and corresponding 95% confidence intervals were calculated. All *P* values were two-sided, with *P* < 0.05 considered statistically significant.

## Results

### Expression and prognostic associations of *RGS10* in breast cancer

To determine the role of RGS10 in breast cancer, we analyzed RGS10 mRNA levels in normal tissues (n = 31) from the Genotype-Tissue Expression dataset, which showed RGS10 mRNA levels were high in normal breast, blood, colon, and small intestine tissues and low in normal heart, liver, and pancreas tissues (Figure 1A). Next, we determined RGS10 mRNA levels in cell lines representing 21 human cancers from the Cancer Cell Line Encyclopedia database (Figure 1B). Finally, we applied RT-qPCR in freshly resected breast cancer tissues (n = 20) and matched adjacent normal breast tissues, and showed RGS10 mRNA levels were lower in breast cancer tissues compared to normal breast tissues (*P* = 0.003; Figure 1C). The clinicopathological characteristics of these 20 patients are shown in Table 1. This pattern implied a downregulation of RGS10 expression in breast cancer tissues.

**Figure 1.**
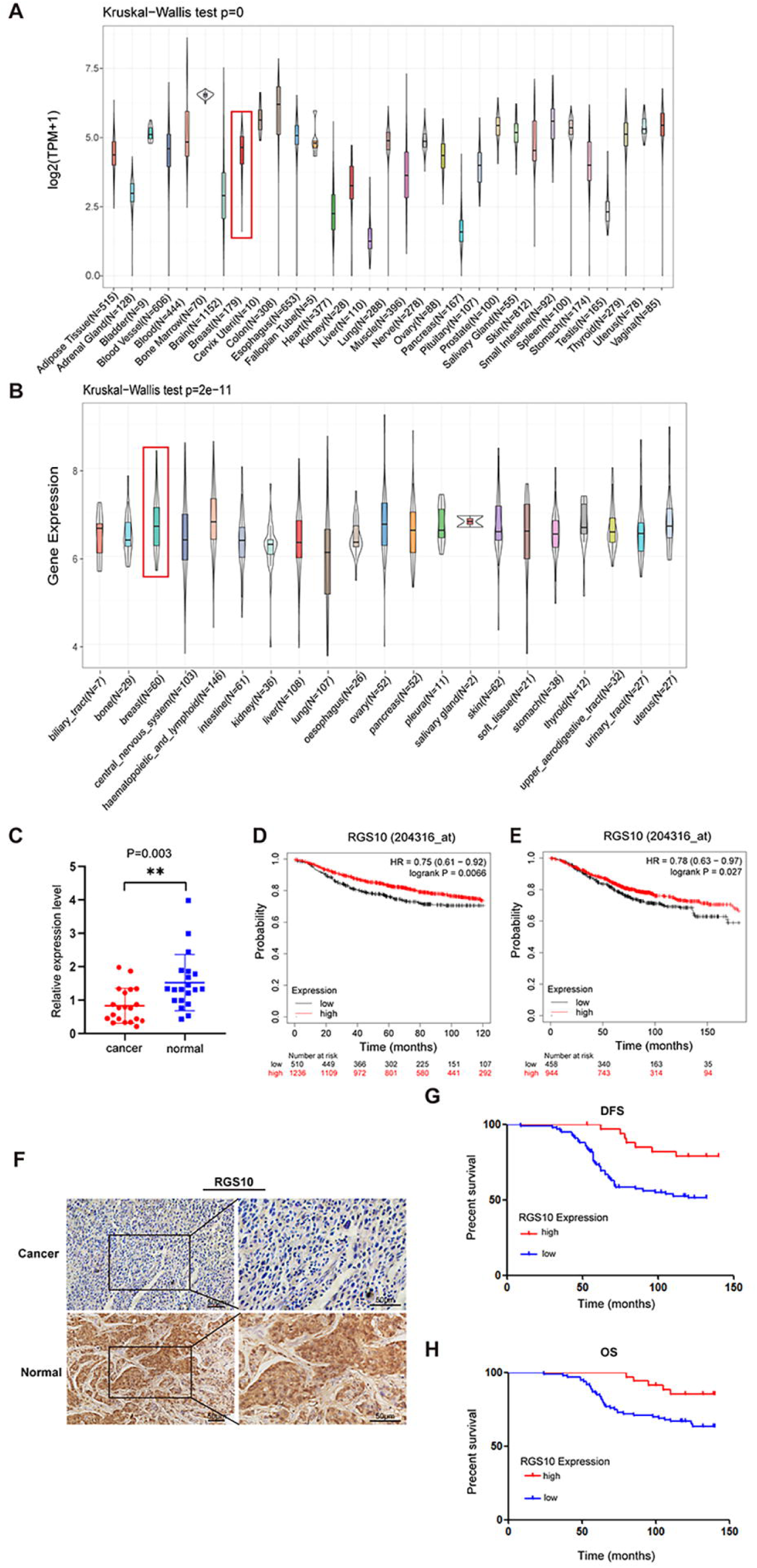
The expression and prognostic associations of RGS10 in breast cancer. (A) RGS10 mRNA levels in 31 normal human tissues. Data were derived from the Genotype-Tissue Expression database. (B) RGS10 mRNA levels in cell lines representing 21 human cancers. Data were derived from the Cancer Cell Line Encyclopedia database. (C) qRT-PCR showing RGS10 mRNA levels in freshly resected breast cancer tissues (n = 20) and matched adjacent normal breast tissues. (D–E) Survival analyses showing disease-free survival (DFS) (D) and overall survival (OS) (E) in patients with breast cancer stratified by high versus low RGS10 mRNA levels. Data were derived from the Kaplan–Meier plotter database. (F) Representative images showing immunohistochemical staining of RGS10 protein expression in breast cancer tissues or normal tissues (n = 133) (magnification: 200× and 400×). (G–H) Kaplan–Meier analysis showing DFS (G) and OS (H) in patients with breast cancer stratified by presence versus absence of RGS10 protein in breast cancer tissues (n = 133).

To investigate the biological role and clinical and prognostic significance of RGS10 in breast cancer tissues, we used survival analyses. In breast cancer samples from the Kaplan–Meier plotter database, high RGS10 mRNA level was associated with significantly improved DFS (*P* = 0.0066; Figure 1D) and OS (*P* = 0.027; Figure 1E). In surgically resected breast cancer tissues (n=133), RGS10 protein expression level detected by immunohistochemistry (representative images shown in Figure 1F) was positively correlated with breast cancer subtype (*P* = 0.043), distant metastasis (*P* = 0.008), and survival status (*P* = 0.024). There were no correlations with age, comorbid disease, histological grade, tumor size, number of positive axillary lymph nodes, number of pregnancies, number of births, age at first pregnancy, menopausal status, estrogen receptor status, progesterone receptor status, human epidermal growth factor receptor 2 (HER2) status, or Ki67 status (Table 2). In these patients, a high RGS10 protein expression level was associated with a longer DFS (*P =* 0.003, Figure 1G) and OS (*P* = 0.022, Figure 1H). Clinicopathological characteristics associated with DFS and OS were identified with Cox regression analyses. On multivariate regression analysis, histological grade and RGS10 protein expression were independent predictors of DFS (Table 3). On univariate regression analysis, age, histological grade, and RGS10 protein expression were independent predictors of OS (Table 4). This suggests low RGS10 expression is associated with poor survival in patients with breast cancer.

**Table 3.**
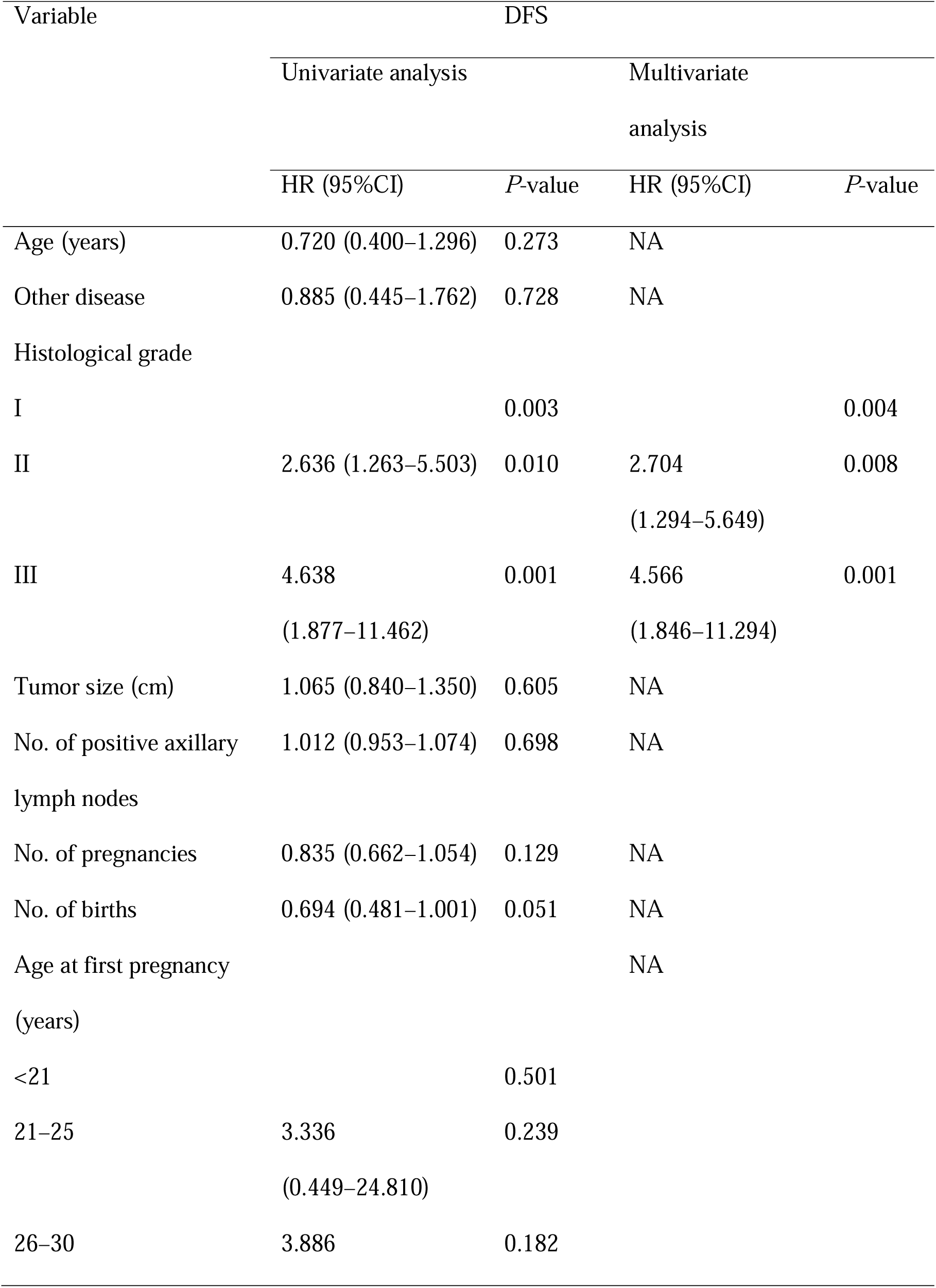
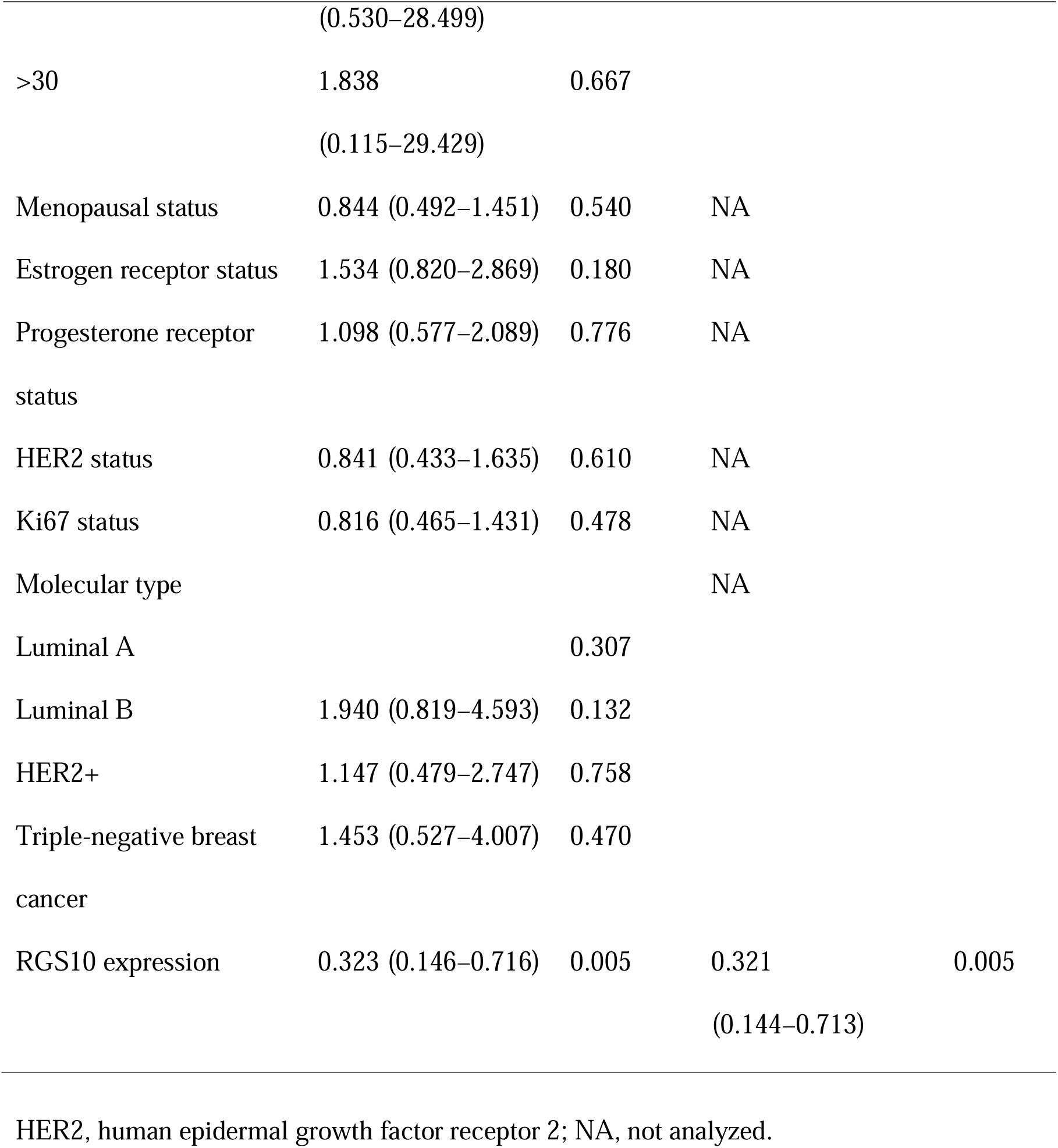
Univariate and multivariate Cox regression analyses of clinicopathological risk factors for disease-free survival.

**Table 4.** Univariate and multivariate Cox regression analyses of clinicopathological risk factors for overall survival.

| Variable | Overall survival |  |  |  |
| --- | --- | --- | --- | --- |
|  | Univariate analysis |  | Multivariate analysis |  |
|  | HR (95%CI) | *P*-value | HR (95%CI) | *P*-value |
| Age (years) | 0.423 (0.226–0.792) | 0.007 | 0.305 (0.154–0.603) | 0.001 |
| Other disease | 0.984 (0.453–2.136) | 0.967 | NA |  |
| Histological grade |  |  |  |  |
| I |  | 0.067 |  | 0.008 |
| II | 1.519 (0.703–3.284) | 0.288 | 1.834 (0.840–4.007) | 0.128 |
| III | 3.062 (1.180–7.943) | 0.021 | 5.090 (1.812–14.300) | 0.002 |
| Tumor size (cm) | 1.045 (0.788–1.385) | 0.760 | NA |  |
| No. of positive axillary lymph nodes | 1.011 (0.943–1.083) | 0.759 | NA |  |
| No. of pregnancies | 0.941 (0.731–1.213) | 0.640 | NA |  |
| No. of births | 0.896 (0.625–1.285) | 0.550 | NA |  |
| Age at first pregnancy (years) |  |  | NA |  |
| <21 |  | 0.580 |  |  |
| 21–25 | 2.818 (0.378–21.001) | 0.312 |  |  |
| 26–30 | 2.021 (0.269–15.190) | 0.494 |  |  |
| >30 | 3.255 (0.295–35.934) | 0.335 |  |  |
| Menopausal status | 0.769 (0.405–1.458) | 0.420 | NA |  |
| Estrogen receptor status | 0.769 (0.405–1.458) | 0.420 | NA |  |
| Progesterone receptor status | 1.091 (0.519–2.293) | 0.818 | NA |  |
| HER2 status | 0.587 (0.246–1.398) | 0.229 | NA |  |
| Ki67 status | 0.718 (0.370–1.392) | 0.327 | NA |  |
| Molecular type |  |  | NA |  |
| Luminal A |  | 0.589 |  |  |
| Luminal B | 0.639 (0.295–1.383) | 0.255 |  |  |
| HER2+ | 0.756 (0.272–2.100) | 0.592 |  |  |
| Triple-negative breast cancer | 1.079 (0.467–2.493) | 0.860 |  |  |
| RGS10 expression | 0.353 (0.138–0.900) | 0.029 | 0.364 (0.154–0.603) | 0.001 |

HER2, human epidermal growth factor receptor 2; NA, not analyzed.

Taken together, these findings imply that RGS10 has a role in suppressing breast cancer and RGS10 may represent a potential prognostic biomarker in breast cancer.

### RGS10 silencing increases the proliferation and migration of breast cancer cells *in vitro*

To characterize RGS10 protein expression in the breast cancer cell lines MDA-MB-231, MCF7, and SKBR3, we conducted Western blotting. MDA*-*MB*-*231 is a highly aggressive, invasive, and poorly differentiated triple-negative breast cancer cell line (Bianchini et al., 2016). MCF7 is an adenocarcinoma cell line with estrogen, progesterone, and glucocorticoid receptors (Lee et al., 2015). The SKBR3 cell line overexpresses the HER2/c-erb-2 gene product (Merlin et al., 2002). RGS10 protein levels appeared lower in the highly aggressive cell line MDA-MB-231 compared to the poorly aggressive and less invasive cell lines MCF7 and SKBR3 (Figure 2A). This finding validates our previous observations suggesting that RGS10 acts as a tumor suppressor in breast cancer.

**Figure 2.**
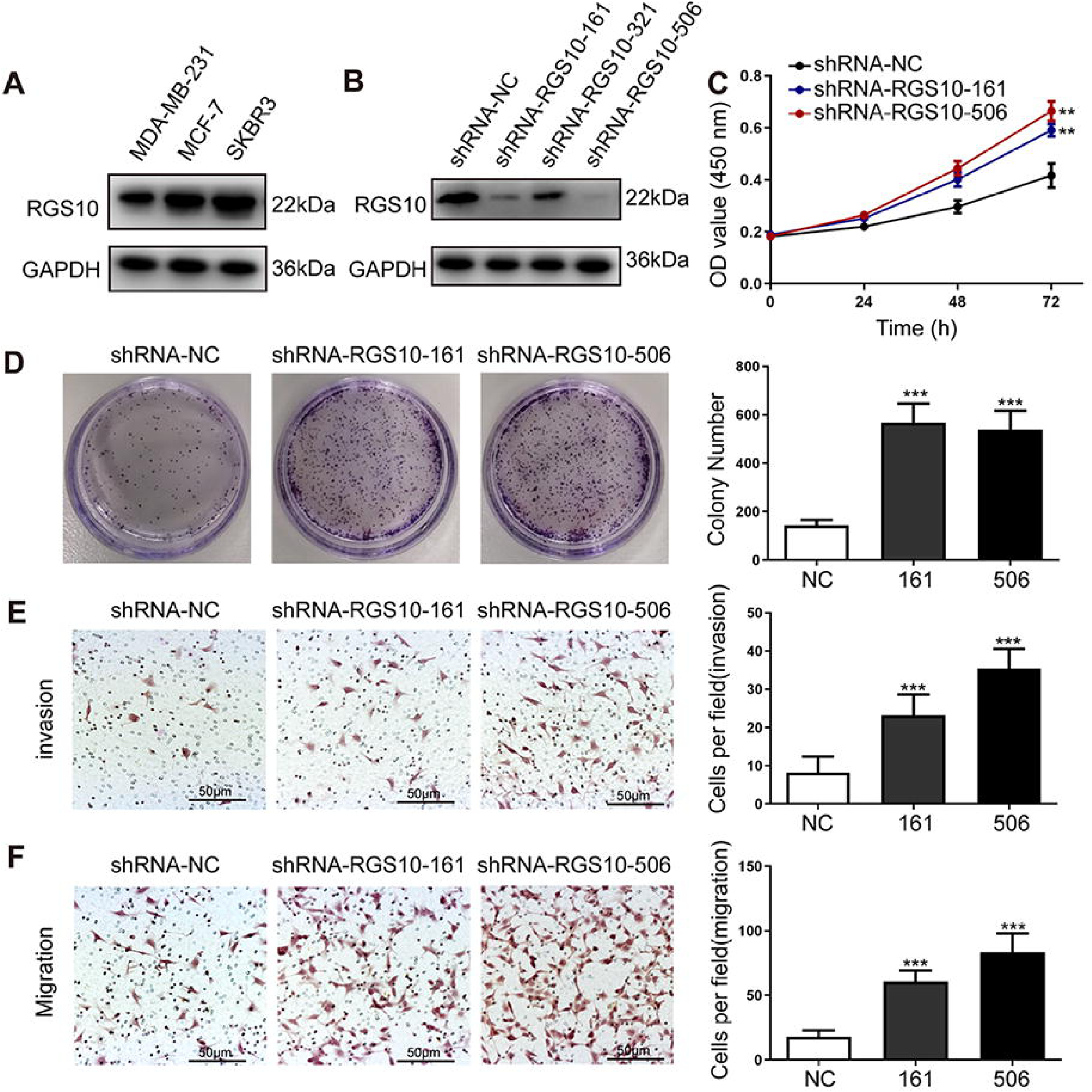
RGS10 silencing increases the proliferation and migration of breast cancer cells *in vitro*. (A) Western blotting showing RGS10 protein levels in molecularly distinct breast cancer cell lines. (B) Western blotting showing RGS10 protein levels in SKBR3 cells transfected with three independent shRNA constructs, shRNA-RGS10-161, shRNA-RGS10-321, and shRNA-RGS10-506, and shRNA-NC. (C) CCK-8 assay showing the viability of SKBR3 cells transfected with shRNA-RGS10-161, shRNA-RGS10-506, or shRNA-NC. (D–F) Colony formation (D) and transwell migration/invasion (E, F) assays in SKBR3 cells transfected with shRNA-RGS10-161, shRNA-RGS10-506, or shRNA-NC.

To investigate the role of RGS10 in breast cancer cells, we silenced RGS10 in SKBR3 cells using two independent shRNA constructs, shRNA-RGS10-161 and shRNA-RGS10-506, which had significantly improved transfection efficiency compared to shRNA-RGS10-321 (Figure 2B). We evaluated the effects of RGS10 on breast cancer cell proliferation, colony formation, migration, and invasion in RGS10-depleted (shRNA-RGS10-506, shRNA-RGS10-161) SKBR3 cells and SKBR3 cells transfected with shRNA-negative control (NC). Cell Counting Kit-8 (CCK-8) analysis, Colony formation, and Transwell migration/invasion assays showed significantly increased proliferative ability, clonogenic ability, migration ability, and invasive capacity in RGS10-depleted SKBR3 cells compared to NC (P < 0.001) (Figure 2C–F).

These findings show invasion and metastasis were enhanced in breast cancer cells lacking RGS10, suggesting an inhibitory effect of RGS10 in breast cancer metastasis.

### RGS10 silencing promotes lipocalin-2 (LCN2) expression and EMT in breast cancer cells

To explore the mechanisms by which RGS10 suppresses breast cancer invasion and metastasis, we analyzed potential downstream tumor metastasis-related genes by comparing the transcriptomes in RGS10-depleted (shRNA-RGS10-506, shRNA-RGS10-161) SKBR3 cells and SKBR3 cells transfected with shRNA-NC. Differential gene expression was visualized using a volcano plot, which revealed that 70 genes were significantly upregulated in RGS10-depleted SKBR3 cells (Figure 3A). KEGG pathway analysis and GO enrichment analysis of differentially expressed genes identified upregulated KEGG pathways, biological processes, cellular components, and molecular functions (Figure 4B–E). Upregulated KEGG pathways were associated with cytokine-cytokine receptor interactions and extracellular matrix-receptor interactions. Gene sets associated with high- and low-RGS10 mRNA expression were identified by gene set enrichment analysis using the Molecular Signatures Database (Liberzon et al., 2011). Gene sets associated with high-RGS10 mRNA expression included allograft rejection, apoptosis, interleukin (IL) 6/Janus kinase/signal transducer and activator of transcription pathway (STAT) 3, IL2/STAT5 pathway, and inflammatory response (Table 5). Western blotting showed changes in biomarkers of EMT in RGS10-depleted SKBR3 cells compared to NC. Neutrophil-derived cytokine LCN2 and vimentin protein levels were higher and E-cadherin protein levels were lower in RGS10-depleted SKBR3 cells compared to NC (Figure 3F–G).

**Figure 3.**
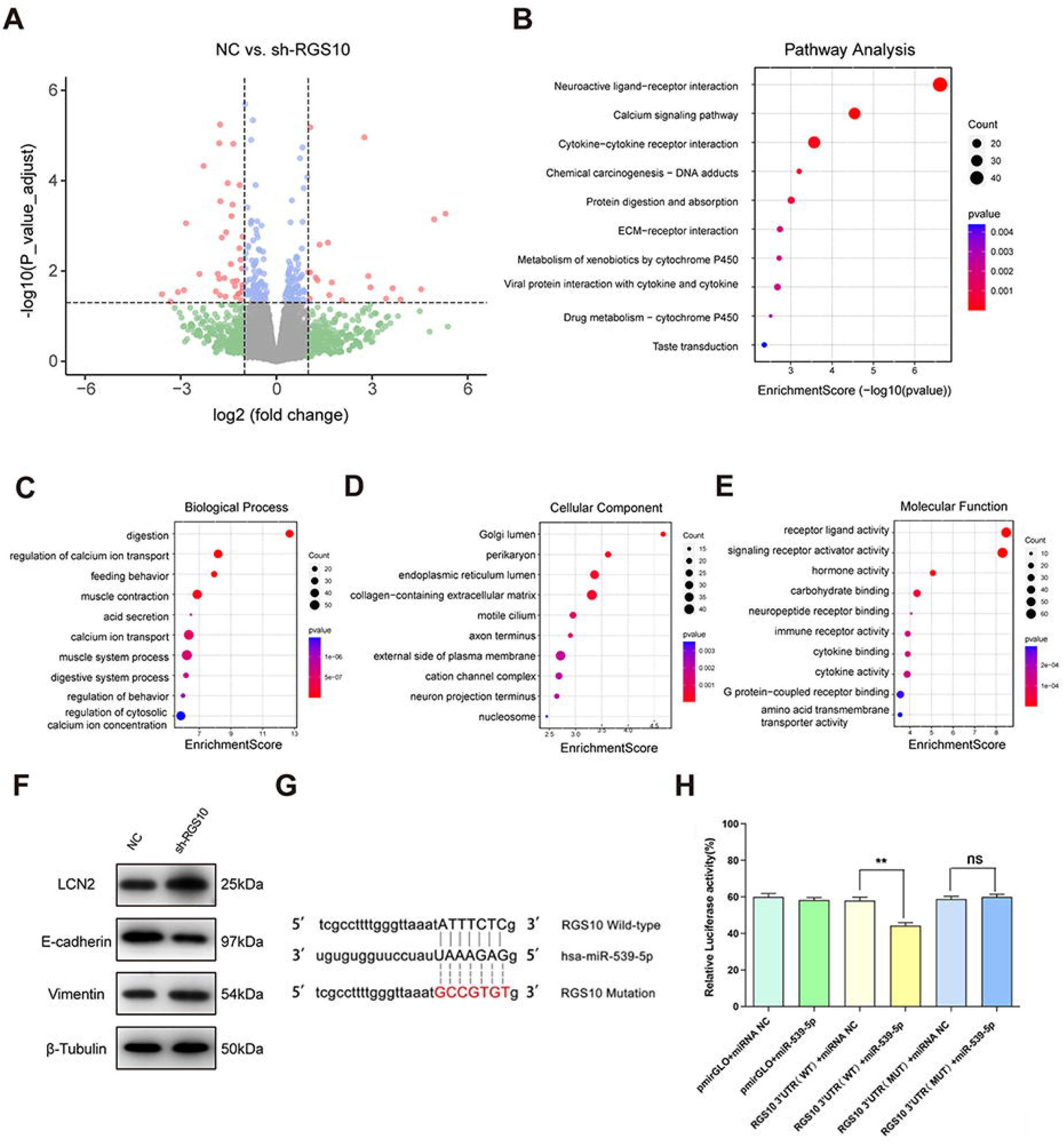
Protein–protein network interaction network and GO functional- and KEGG pathway-enrichment analysis of genes co-expressed with RGS10. (A) Volcano plot showing differentially expressed genes between SKBR3 cells transfected with shRNA-RGS10 or shRNA-NC. (B–E) KEGG pathway analysis and GO enrichment analysis of differentially expressed genes showing the ten most enriched terms. BP: biological processes; MF: molecular function; CC: cellular compartment. (F) LCN2, E-cadherin, and vimentin protein levels in SKBR3 cells transfected with shRNA-RGS10 or shRNA-NC. (G) Schematics of predicted miR-539-5p binding sites between wild-type and mutant RGS10 sequences in the 3′-untranslated regions. (H) Relative luciferase activities detected after cotransfection of wild-type or mutant luciferase reporter plasmids and an miR-539-mimic.

**Figure 4.**
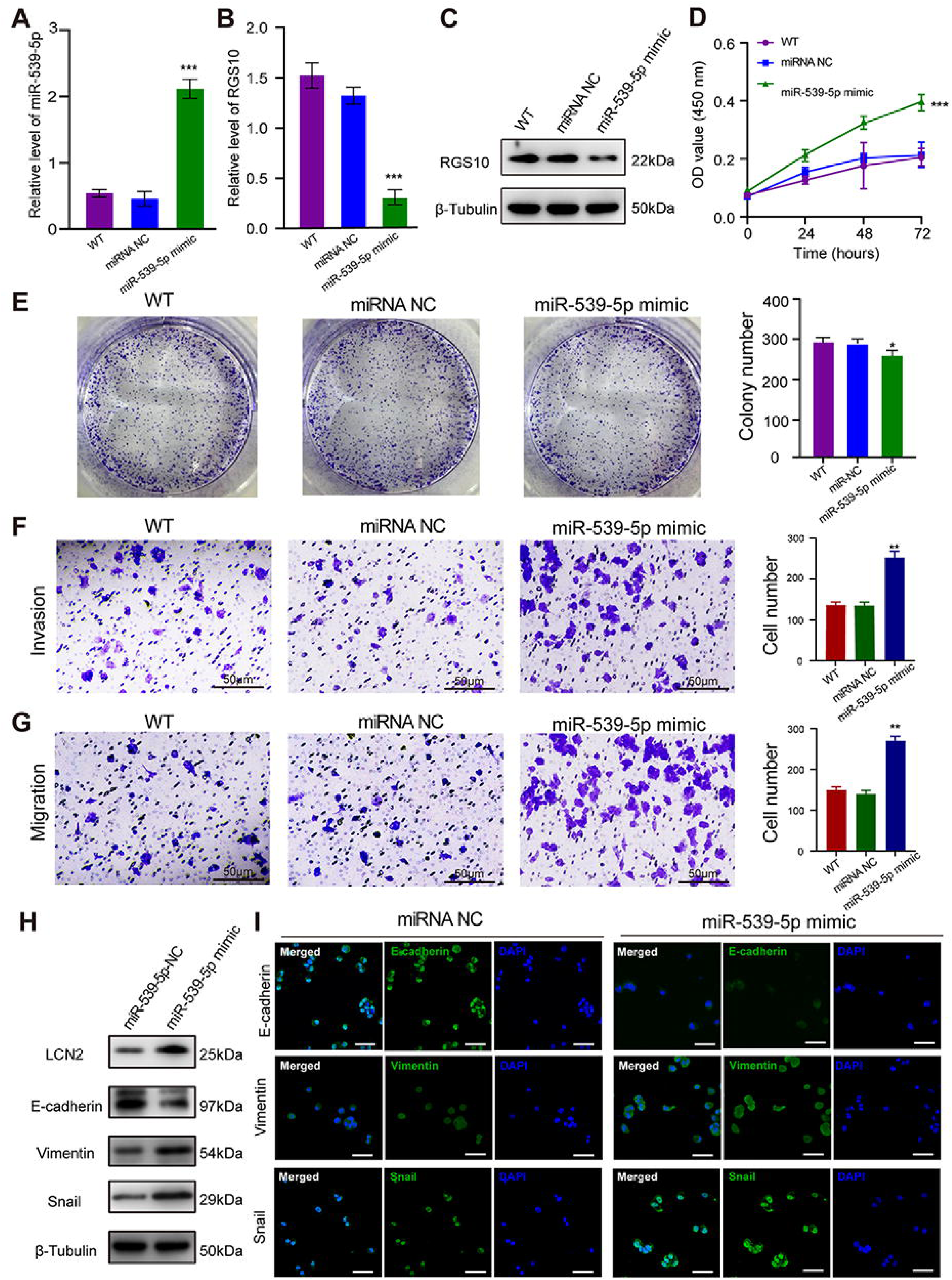
miR-539-5p regulates the migration, invasion, proliferation, and EMT of breast cancer cells. (A) qPCR showing the transfection efficiency of the miR-539-5p mimic. *** *P* < 0.001, one-way ANOVA. (B–C) qRT-PCR and Western blotting showing RGS10 mRNA and protein levels in SKBR3 cells transfected with the miR-539-5p mimic, negative control (NC), or wild type (WT). *** *P* < 0.001, one-way ANOVA. (D) CCK-8 assay showing the viability of SKBR3 cells transfected with the miR-539-5p mimic, NC, or WT. *** *P* < 0.001, one-way ANOVA. (E–G) Colony formation (E) and transwell migration/invasion (F, G) assays in SKBR3 cells transfected with the miR-539-5p mimic, NC, or WT. * *P* < 0.05, ** *P* < 0.01, Student’s t test. (H) Western blotting showing protein levels of LCN2 and biomarkers of EMT in SKBR3 cells transfected with the miR-539-5p mimic or NC. (I) Immunofluorescence staining showing E-cadherin, vimentin, and snail protein expression in SKBR3 cells transfected with the miR-539-5p mimic or NC. Scale bar: 50 µm.

**Table 5.**
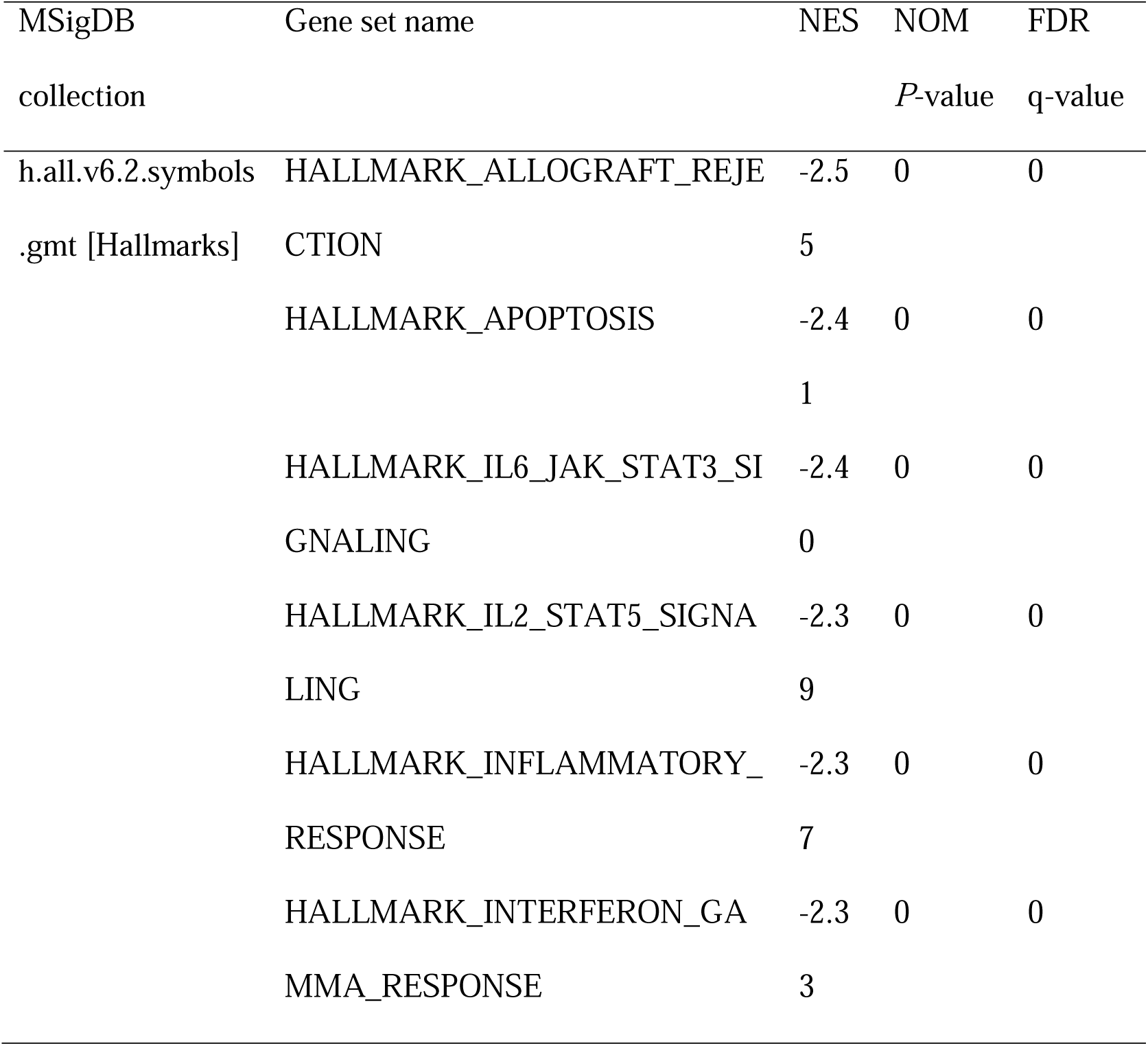
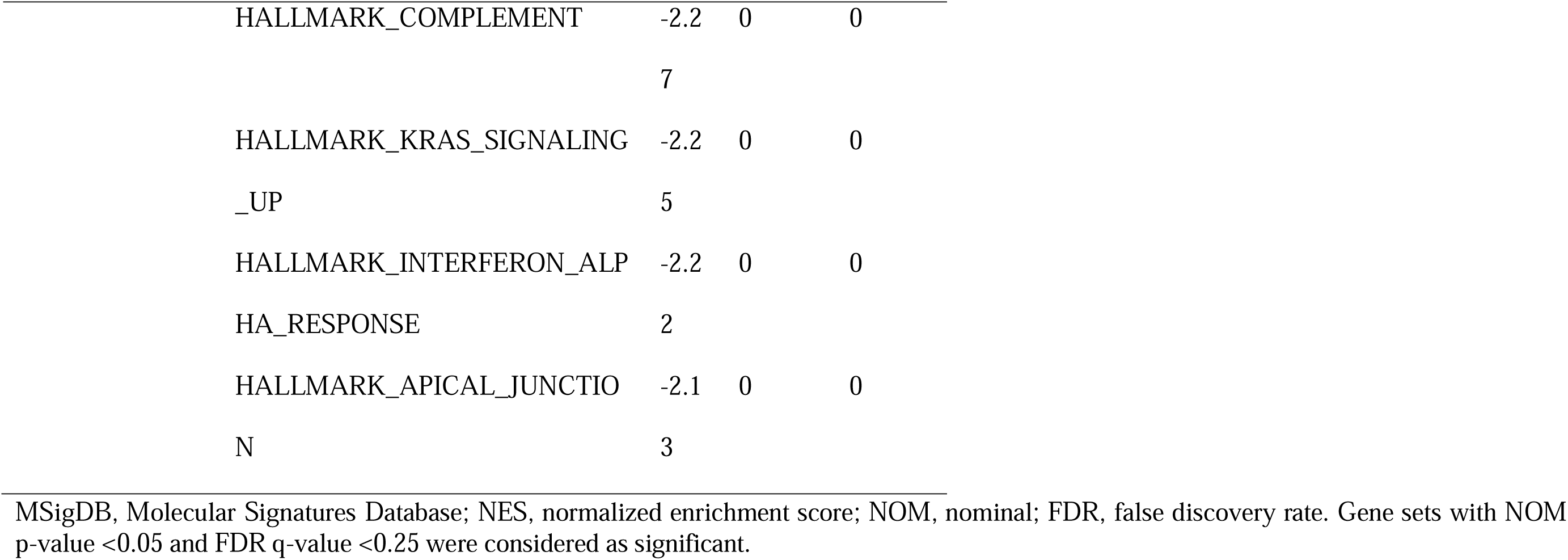
Gene sets enriched in phenotype ‘High’.

These findings show that RGS10 deficiency promotes invasion and metastasis by activating the LCN2 pathway to induce EMT in breast cancer cells, supporting the potential of RGS10 as a prognostic biomarker in breast cancer.

### miR-539-5p regulates the migration, invasion, proliferation, and EMT of breast cancer cells

To study the upstream regulatory mechanism of RGS10 in breast cancer, we used the StarBase database to predict miRNAs that could potentially bind to RGS10. The luciferase reporter assay identified miR-539-5p as a miRNA that targets RGS10 in breast cancer cells (Figure 3G–H).

To predict the potential effects of miR-539-5p on breast cancer cells, we transfected SKBR3 and MDA-MB-231 cells with a miR-539-5p mimic to represent miR-539-5p overexpression, a miR-539-5p inhibitor, or appropriate NCs (Figure 4A). RT-qPCR and Western blotting validated transfection efficiency and showed that RGS10 mRNA and protein levels were significantly decreased in SKBR3 cells overexpressing miR-539-5p compared to SKBR3 cells transfected with miRNA-NC or the wild type (Figure 4B, C). Cell Counting Kit-8 (CCK-8) analysis, and colony formation and Transwell migration/invasion assays showed SKBR3 cell proliferation, colony formation, migration, and invasion were significantly increased in SKBR3 cells overexpressing miR-539-5p compared to SKBR3 cells transfected with miRNA-NC or the wild type (Figure 4D–G). In contrast, RGS10 protein levels were significantly increased (Figure 5A–C) and MDA-MB-231 cell proliferation, colony formation, migration, and invasion were significantly decreased in MDA-MB-231 cells transfected with miR-539-5p inhibitor compared to anti-miRNA-NC or the wild type (Figure 5D–G).

**Figure 5.**
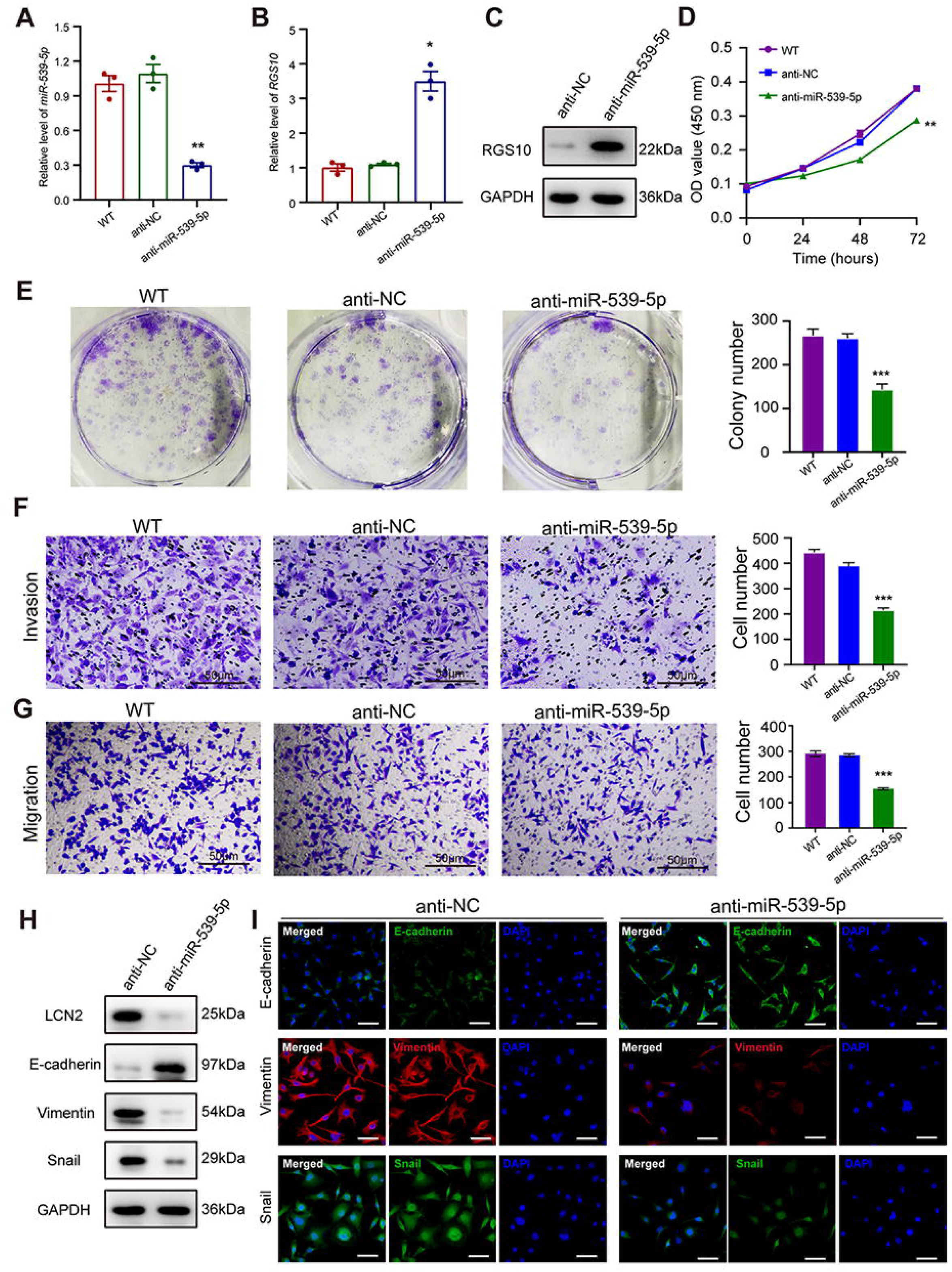
miR-539-5p inhibitor suppresses breast cancer cell proliferation and invasion. (A) qPCR showing transfection efficiency of the miR-539-5p inhibitor after 48 h. *** *P* < 0.001, one-way ANOVA. (B–C) qRT-PCR and Western blotting showing RGS10 mRNA and protein levels in MDA-MB-231 cells transfected with the miR-539-5p inhibitor, negative control (NC), or wild type (WT). *** *P* < 0.001, one-way ANOVA. (D) CCK-8 assay showing the viability of MDA-MB-231 cells transfected with the miR-539-5p inhibitor, NC, or WT. *** *P* < 0.001, one-way ANOVA. (E–G) Colony formation (E) and transwell migration/invasion (F, G) assays in MDA-MB-231 cells transfected with the miR-539-5p inhibitor, NC, or WT. * *P* < 0.05, ** *P* < 0.01, Student’s t test. (H) Western blotting showing protein levels of LCN2 and biomarkers of EMT in MDA-MB-231 cells transfected with the miR-539-5p inhibitor or NC. (I) Immunofluorescence staining showing E-cadherin, vimentin, and snail protein expression in MDA-MB-231 cells transfected with the miR-539-5p inhibitor or NC. Scale bar: 50 µm.

Based on these findings, for the first time, we propose a miR-539-5p/RGS10/LCN2 regulatory axis in breast cancer. Consistent with this, Western blotting and immunofluorescence assays showed decreased E-cadherin protein expression and increased LCN2, vimentin, and snail protein expression in SKBR3 cells overexpressing miR-539-5p compared to SKBR3 cells transfected with miRNA-NC or the wild type (Figure 4H–I). These effects were reversed in MDA-MB-231 cells transfected with a miR-539-5p inhibitor compared to anti-miRNA-NC or the wild type (Figure 5H–I).

These findings identify miR-539-5p as a critical factor in breast cancer metastasis by regulating RGS10/LCN2 expression.

### RGS10 inhibits breast cancer growth by targeting LCN2 *in vivo*

To investigate tumorgenicity, we subcutaneously transplanted SKBR3 cells transfected with shRNA-RGS10-506 or shRNA-NC into nude mice. The growth of tumors derived from RGS10-depleted SKBR3 cells was significantly increased compared to NC, manifested by larger tumor size compared to NC (Figure 6A–C). Immunohistochemistry showed LCN2, snail, and vimentin protein expression were increased and E-cadherin protein expression was decreased in tumor tissues derived from RGS10-depleted SKBR3 cells compared to NC (Figure 6D).

**Figure 6.**
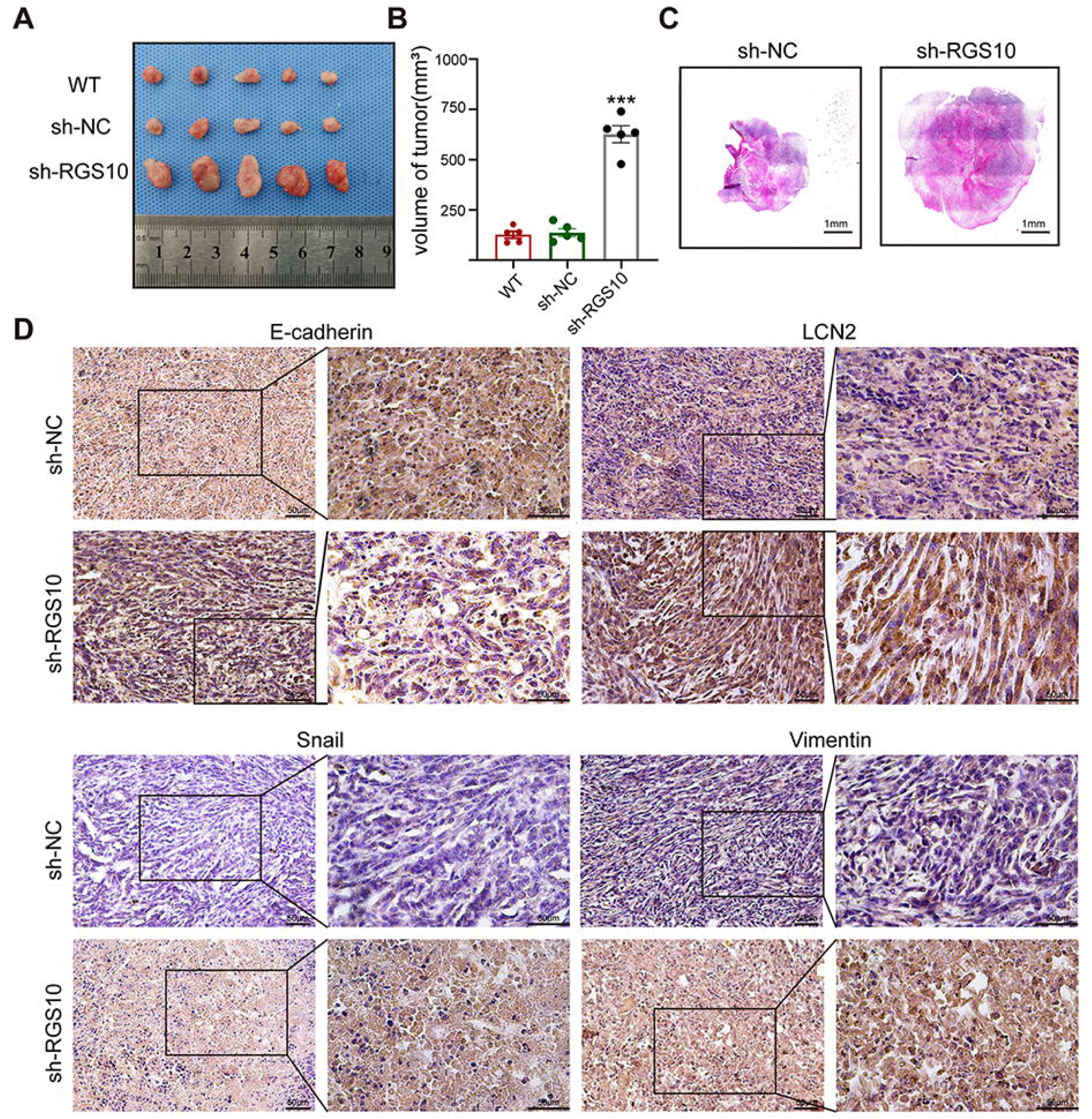
RGS10 inhibits breast cancer growth by targeting LCN2 *in vivo*. (A) Size of tumors derived from RGS10-depleted SKBR3 cells, negative control (NC) and wild type(WT). (B) Volume of tumors derived from RGS10-depleted SKBR3 cells, NC and WT. (C) Hematoxylin and eosin staining of tumors derived from RGS10-depleted SKBR3 cells and NC. (D) Immunohistochemical staining showing LCN2, E-cadherin, snail, and vimentin protein expression in tumors derived from RGS10-depleted SKBR3 cells and NC.

These *in-vivo* data confirm the previous *in-vitro* observations that RGS10 deficiency promotes invasion and metastasis by activating the LCN2 pathway to induce EMT in breast cancer cells, and demonstrate that RGS10 has utility as a prognostic biomarker in breast cancer.

## Discussion

Distant metastasis is a main cause of death in patients with breast cancer. Identification of prognostic biomarkers for early distant metastasis may inform clinical decision-making and improve patient outcomes. To the best of our knowledge, this is the first study to characterize the role of RGS10 as a tumor suppressor and biomarker of EMT in breast cancer. RGS10 was expressed at lower levels in breast cancer tissues than in adjacent normal breast tissues. RGS10 expression was associated with molecular subtype of breast cancer, distant metastasis, and survival status. Patients with high compared to low RGS10 mRNA expression in breast cancer tissues had improved DFS and OS. RGS10 protein levels were lower in the highly aggressive breast cancer cell line MDA-MB-231 compared to the poorly aggressive and less invasive breast cancer cell lines MCF7 and SKBR3. RGS10 reduced breast cancer cell proliferation, colony formation, invasion, and migration by inhibiting EMT via a novel mechanism dependent on LCN2 and miR-539-5p.

EMT is involved in normal development and morphogenic processes, including embryogenesis and tissue regeneration. Pathological EMT promotes invasion and metastasis in tumors through intracellular signaling, transcription factors, miRNAs, and epigenetic and posttranslational regulators (Huang et al., 2022; Lamouille et al., 2014). Signaling pathways such as TGF-β, Wnt, Notch, and phosphoinositide 3-kinase/AKT contribute to EMT, often with cross-talk at various levels and feedback activation/repression mechanisms (Deshmukh et al., 2021; Huang et al., 2022). Transcription factors such as snail, slug, ZEBl/ZEB2, and Twist1/Twist2 induce EMT by acting on the E-box sequence of the CDH1 promoter. Noncoding miRNAs regulate EMT by selectively targeting mRNAs of cell receptors, signaling pathways, the cell cycle, or cell adhesion (Górecki and Rak, 2021; Huang et al., 2022; Li et al., 2019). Epigenetic modifications, including DNA methylation and histone modifications, alter the expression of EMT transcription factors involved in the molecular pathways of metabolism, transcription, differentiation, and apoptosis (Górecki and Rak, 2021).

The present study identifies RGS10 as an important mediator of EMT in breast cancer. Previous studies have demonstrated links between several RGS proteins and various cancers, with RGS proteins acting as tumor initiators or suppressors depending on the RGS protein and type of cancer (Li et al., 2023). RGS10 is the smallest protein of the RGS D/12 subfamily, which functions as GAPs for the Gi family Gα subunits (Almutairi et al., 2020). RGS10 has been linked to a poor prognosis in patients with laryngeal cancer (Yin et al., 2013), liver cancer (Wen et al., 2015), and childhood acute myeloid leukemia (Chaudhury et al., 2018). RGS10 may represent a biomarker of clinical staging for ovarian cancer and is one of five signature genes involved in the occurrence and development of ovarian cancer. This five-gene signature (RGS11, RGS10, RGS13, RGS4, and RGS3) is overexpressed in ovarian cancer and involved in extracellular matrix-receptor interaction, the TGF-β signaling pathway, the Wnt signaling pathway, and the chemokine signaling pathway. These pathways mediate the proliferation, migration, and invasion of ovarian cancer cells; in particular, TGF-β signaling plays an important role in EMT in ovarian cancer (Hu et al., 2021).

The novel action of RGS10 in EMT in breast cancer appears to be dependent on LNC2, also known as neutrophil gelatinase-associated lipocalin, siderocalin, uterocalin, and oncogene 24p3. LNC2 is a secreted glycoprotein of the adipokine superfamily. LCN2 expression levels are particularly high in breast, pancreatic, ovarian, colorectal, thyroid, and bile duct cancer tissues and tumor cell lines. Previous studies show LCN2 can promote tumorigenesis by increasing invasion, metastasis, and proliferation while decreasing apoptosis, possibly because LCN2 can facilitate iron intake to cancer cells and form a heterodimer with matrix metalloproteinase-9 (Santiago-Sánchez et al., 2020). In breast cancer, LCN2 can promote progression by inducing EMT through the estrogen receptor alpha/Slug axis (Morales-Valencia et al., 2022; Yang et al., 2009).

The regulation of EMT in breast cancer by RGS10 may rely on upstream regulation by miR-539-5p. miRNAs play a critical role in the cellular processes of breast cancer, including EMT (Zhang et al., 2013; Zhao et al., 2017). Previous reports indicate that miR-539 expression is downregulated in breast cancer tissues and cell lines (Guo et al., 2018), and miR-539 acts as a tumor suppressor by targeting epidermal growth factor receptor (Guo et al., 2018), specificity protein 1 (Cai et al., 2020), or laminin subunit alpha 4 (Cai et al., 2020) expression. In patients with breast cancer, decreased expression of miR-539 was significantly associated with lymph node metastasis. In breast cancer cells, overexpression of miR-539 inhibited the proliferation and promoted apoptosis of breast cancer cells, suppressed EMT, and sensitized cells to cisplatin treatment (Cai et al., 2020). Further studies are required to fully elucidate miRNA regulation of the gene expression networks that are essential to EMT in breast cancer.

Inflammation promotes EMT in tumors, and EMT induces the production of proinflammatory factors by cancer cells (Suarez-Carmona et al., 2017). The present study showed that RGS10 expression in SKBR3 cells was associated with the inflammatory response. Previous reports show that RGS10 regulates cellular physiology and fundamental signaling pathways in microglia, macrophages, and T-lymphocytes. In microglia and ovarian cancer cells, RGS10 regulates inflammatory signaling by a G protein-independent mechanism, linking RGS10 to the inflammatory signaling mediators tumor necrosis factor-alpha (TNFα) and cyclooxygenase-2 (Alqinyah et al., 2018). In macrophages, RGS10 regulates activation and polarization by suppressing the production of the inflammatory cytokines TNFα and IL6 (Dean and Hooks, 2020). These findings suggest a potential role for RGS10 in the development of an inflamed and immunosuppressive tumor microenvironment, which may have important implications for the progression in breast cancer (Suarez-Carmona et al., 2017).

Our study had some limitations. First, this was a retrospective study with a small sample size. Thus, the conclusions should be confirmed in meta-analyses or large randomized controlled trials. Second, the mechanism by which the miR-539-5p/RGS10/LCN2 axis may be related to outcomes in patients with breast cancer remains to be elucidated. Biochemical characterization of the molecular mechanisms of RGS10 in breast cancer should provide additional insight into the potential of RGS10 as a biomarker of EMT, metastasis, and prognosis in breast cancer and the role of RGS10 as a therapeutic target.

In conclusion, the results of this study show that RGS10 expression is related to survival outcomes in patients with breast cancer. RGS10 protein levels were lower in the highly aggressive breast cancer cell line MDA-MB-231 compared to the poorly aggressive and less invasive breast cancer cell lines MCF7 and SKBR3. Silencing RGS10 expression effectively increased the proliferation, colony formation, invasion, and migration ability of SKBR3 cells. The miR-539-5p/RGS10/LCN2 pathway was identified as an important regulatory axis of EMT in breast cancer. These data demonstrate that RGS10 may play a tumor suppressor role and be considered a biomarker of EMT and prognosis in breast cancer.

## Declarations

### Ethics approval and consent to participate

This study was approved by the Institutional Review Board of Shengjing Hospital of China Medical University.

### Consent for publication

Not applicable

### Availability of data and material

These data and materials are available from the corresponding author for rational reasons.

### Authors’ contributions

Xi Gu and Yang Liu designed this research. Yi Jiang and Yang Liu analyzed the data derived from public databases. Yang Liu, Yi Jiang, Peng Qiu, Tie Ma, Yang Bai, Tong Zhu, Jiawen Bu, Yueting Hu, and Ming Jin performed the experiments. Xi Gu and Yi Jiang wrote the manuscript. All authors read and approved the final manuscript.

